# Temporo-Parietal Cortex Involved in Modeling One’s Own and Others’ Attention

**DOI:** 10.1101/2020.10.04.325357

**Authors:** Arvid Guterstam, Branden J Bio, Andrew I Wilterson, Michael SA Graziano

**Affiliations:** Department of Psychology, Princeton University, Princeton, NJ 08544

## Abstract

In a traditional view, in social cognition, attention is equated with gaze and people track attention by tracking other people’s gaze. Here we used fMRI to test whether the brain represents attention in a richer manner. People read stories describing an agent (either oneself or someone else) directing attention to an object in one of two ways: either internally directed (endogenous) or externally induced (exogenous). We used multivoxel pattern analysis to examine how brain areas within the theory-of-mind network encoded attention type and agent type. Brain activity patterns in the left temporo-parietal junction (TPJ) showed significant decoding of information about endogenous versus exogenous attention. The left TPJ, left superior temporal sulcus (STS), precuneus, and medial prefrontal cortex (MPFC) significantly decoded agent type (self versus other). These findings show that the brain constructs a rich model of one’s own and others’ attentional state, possibly aiding theory of mind.

**Impact statement:** This study used fMRI to show that the human brain encodes other people’s attention in enough richness to distinguish whether that attention was directed exogenously (stimulus-driven) or endogenously (internally driven).

## Introduction

Reconstructing someone else’s attentional state is of central importance in theory of mind (Baron-Cohen, 1997; Calder et al., 2002; Graziano, 2013). By identifying the object of someone else’s attention, and having some intuitive understanding of the complex dynamics and consequences of attention, one can reconstruct at least some of the other person’s likely thoughts, intentions, and emotions, and make predictions about that person’s behavior. Almost all work on how people reconstruct the attention of others has focused on gaze direction. For example, the human eye has a high contrast between pupil and sclera, possibly an adaptation for better gaze tracking (Kobayashi and Kohshima, 1997). The superior temporal sulcus in monkeys and humans may contain specialized neural circuitry for processing gaze direction (Hoffman and Haxby, 2000; Perrett et al., 1985; Puce et al., 1998; Wicker et al., 1998). Seeing a face gaze at an object automatically draws one’s own attention to the object (Friesen and Kingstone, 1998; Frischen et al., 2007). These and other findings show the importance of reconstructing gaze direction in social cognition.

To be adaptive in aiding theory of mind, however, a model of attention should be far more than a vector indicating gaze direction. We previously suggested that the human brain constructs a rich, dynamic, and predictive model of other people’s attention (Graziano, 2019, 2013; Graziano and Kastner, 2011). The model should contain information about different types of attention, about the rapidity or sluggishness with which attention tends to move from item to item, about how external factors such as salience and clutter are likely to affect a person’s attention, and about how attention profoundly affects thought, memory, and behavior. In the proposal, that deeper model is constrained by incoming information, including gaze direction. However, other cues can also constrain the model. People rely on the other person’s body posture, on cues in the surrounding environment, on speech, and on social context. For example, blind people must be able to build models of other people’s attention without seeing the other person’s eyes. Likewise, during a phone conversation, we cannot see the other person and yet we intuitively understand whether that person is attending to what we have said or is distracted by her own words or by a salient event on her end of the line.

Several recent experiments provide evidence for an automatically constructed model of the attention of others that may go beyond merely registering gaze direction (Guterstam et al., 2019; Guterstam and Graziano, 2020; Kelly et al., 2014; Pesquita et al., 2016; Vernet et al., 2019). For example, Pesquita et al. (2016) found that when participants watch an actor in a video attending to an object, the participants implicitly distinguish between whether the actor’s attention was drawn to the object exogenously (bottom-up, or stimulus-driven attention), or whether the actor endogenously shifted attention to the object (top-down, or internally driven attention). The ability to distinguish between someone else’s exogenous and endogenous attention is one example of how people may construct a rich, dynamic model of other people’s attention beyond merely encoding gaze direction or identifying the object of attention.

Inspired by the vignette-style tasks widely used in studies on theory of mind (Fletcher et al., 1995; Gallagher et al., 2000; Happé, 1994; Saxe and Kanwisher, 2003; Vogeley et al., 2001), in the present study, we used functional magnetic resonance imaging (fMRI) and multi-voxel pattern analysis (MVPA) to study brain activity in participants while they read brief stories about people’s attention. Some of the stories implied that attention was being attracted exogenously (“Kevin walks into his closet and notices the bright red tie…”) and some stories implied that attention was being directed endogenously (“Kevin walks into his closet and looks for the bright red tie…”). We also included analogous stories written in the first person, casting the subject of the experiment as the agent (“You walk into your closet and notice the bright red tie…”). The study therefore used a 2×2 design (exogenous versus endogenous attention X self agent versus other agent). Finally, we included a fifth, control condition, consisting of nonsocial stories in which the agent was replaced by an inanimate object, that, like attention, has a source and a target, such as a camera or a light source (“In a closet, a light shines on a red tie…”).

We made four predictions. Our first, central prediction was inspired by the Pesquita et al. (2016) study described above. We hypothesized that participants would encode the type of attention in the story (exogenous versus endogenous), and that this encoding would be evident in some subset of the areas classically involved in theory of mind. Previous experiments on theory of mind typically recruited a network of cortical areas including the temporoparietal junction (TPJ), the superior temporal sulcus (STS), the medial prefrontal cortex (MPFC), and the precuneus (Gallagher et al., 2000; Saxe and Kanwisher, 2003; van Veluw and Chance, 2014; Vogeley et al., 2001). We therefore predicted that the exogenous-versus-endogenous distinction would be significantly encoded in some subset of these areas. In particular, we anticipated that the TPJ might show the clearest evidence of encoding information about the type of attention, since previous experiments pointed to the TPJ as contributing to encoding other people’s attentional state when participants looked at faces gazing toward objects (Igelström et al., 2016; Kelly et al., 2014). This first prediction, that the social cognition network will encode the exogenous-versus-endogenous distinction, represents the main, novel contribution of this study.

Second, we predicted that participants would encode information about the agent in the story (self versus other), and that this encoding would again be evident in some subset of the areas classically involved in theory of mind. Self-versus-other encoding has been examined in previous studies, and found to be reflected in the theory-of-mind network (e.g., Northoff et al., 2006; Ochsner et al., 2004; Passingham et al., 2010; Qin and Northoff, 2011; van Veluw and Chance, 2014). This second prediction represents a test of whether our present paradigm, using subtle wording differences between similar sentences, can produce results consistent with previous findings.

Third, we predicted that participants would encode information associated with the interaction between the two factors. We predicted that at least some subset of the areas in the theory-of-mind network may encode the type of attention (exogenous versus endogenous) to a different extent in self-related stories as compared to other-related stories.

Fourth and finally, we tested for brain regions that encoded the distinction between social stories (with human agents) and nonsocial stories (with only non-agent objects). We predicted that this social-versus-nonsocial encoding would again be evident in the same network of brain regions noted above, that are known to be involved in theory of mind. This final analysis served as a control to check on the validity of the story stimuli and confirm that they engaged social cognition as expected.

## Methods

### Subjects

Thirty-two healthy human volunteers (12 females, 30 righthanded, aged 18-52, normal or corrected to normal vision) participated in the study, based on the sample size used in a previous, related study (Guterstam et al., 2020). Subjects were recruited either from a paid subject pool, receiving 40 USD for participation, or from among Princeton undergraduate students, who received course credits as compensation. In the subject recruitment material, the experiment was described as a “Reading Comprehension Study.” All subjects provided informed consent and all procedures were approved by the Princeton Institutional Review Board.

### Experimental setup

Before scanning, subjects were instructed and then shown three sample trials (which were not part of the stories presented in the subsequent experiment) on a laptop computer screen. All subjects gave the correct response to all three trials on the first try, indicating they had understood the instructions adequately. During scanning, the subjects laid comfortably in a supine position on the MRI bed. Through an angled mirror mounted on top of the head coil, they viewed a translucent screen approximately 80 cm from the eyes, on which visual stimuli were projected with a Hyperion MRI Digital Projection System (Psychology Software Tools, Sharpsburg, PA, USA) with a resolution of 1920 x 1080 pixels. A PC running MATLAB (MathWorks, Natick, MA, USA) and the Psychophysics Toolbox (Brainard, 1997) was used to present visual stimuli. A right hand 5-button response unit (Psychology Software Tools Celeritas, Sharpsburg, PA, USA) was strapped to the subjects’ right wrist. Subjects used the right index finger button to indicate a true response, and the right middle finger to indicate a false response during the probe phase of each trial.

### Experimental conditions and stimuli

Five experimental conditions were included. Subjects were presented with short stories (2-3 sentences, average word count = 24) describing a scene in which an agent, which was either the subject him-/herself (self) or another person (other), directed attention to something in the external world endogenously (e.g., “X is attentively looking for Y”) or exogenously (e.g., “X’s attention is captured by Y”). These four conditions made up a 2 x 2 factorial design: attention type (endogenous versus exogenous) X agent (self versus other). In addition, we included a baseline condition featuring stories in which the agent was substituted by a non-human object. In each trial, after a 9 – 11 s inter-trial interval, the story was presented for 10 s in easily readable, white text on a black background, at the center of the screen, after which a probe statement was shown for 4 s, to which the subjects responded either true or false by button press. See Figure 1 for details, and SI Data S1 for all stories.

**Figure 1.**
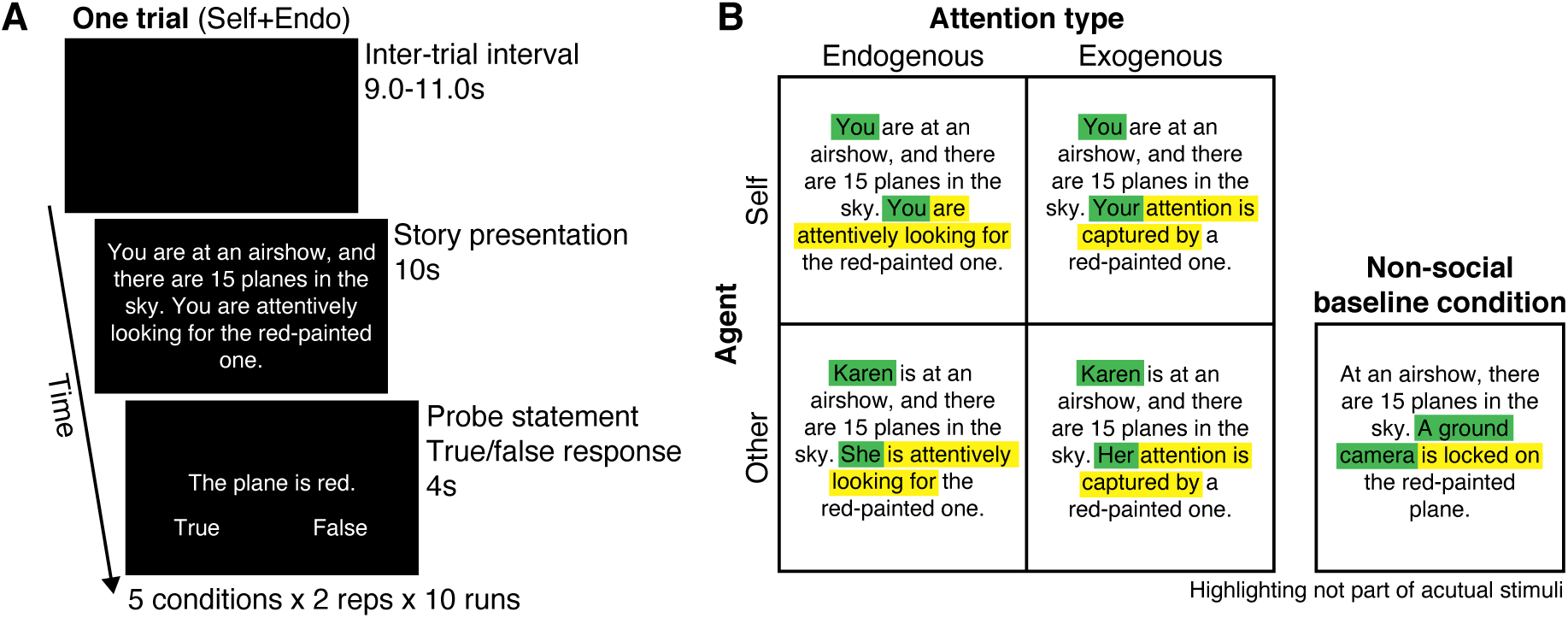
Methods. **A**. Schematic timeline of the fMRI design. In each trial, subjects were presented with a short story for 10 s, describing a scene in which an agent attended to an object in the environment. A probe statement was then shown for 4 s, relating to either the story’s spatial context or object property, to which the subjects responded either true or false by button press. **B**. The agent in the story was either the subject him-/herself (self) or another person (other), and directed attention to the object endogenously (internally driven attention) or exogenously (stimulus-driven attention), yielding a 2 × 2 factorial design of attention type × agent. We created 80 unique stories in four different versions, one for each condition. We made minimal changes to the wordings to keep the story versions as semantically similar as possible. Green highlighting indicates wording specifying agent, yellow highlighting indicates wording specifying attention type (colors not part of actual visual stimuli). For each story, each subject saw only one of the four versions (balanced across subjects). We also included a nonsocial baseline condition (twenty unique stories based on a subset of the 80 social stories) in which the agent was replaced by a non-human object.

Each subject ran 100 trials and thus saw 100 stories: 80 social stories and 20 non-social control stories. The 80 social stories were constructed as follows. We began with 80 unique short stories. For each story, four versions were constructed, one for each of the factorial conditions (Figure 1B). To keep the story versions as semantically similar as possible, we made minimal changes to the wordings. To distinguish the self and other versions, we substituted the word “you” with a name (e.g., “Karen”) and the word “your” with “his” or “her”. The names in the stories were selected from a list of the 100 most popular given names for male and female babies born during the years 1919-2018 in the United States, which is published by the Social Security Administration (https://www.ssa.gov/oact/babynames/decades/century.html). Half of the names were masculine, half feminine. To distinguish the endogenous and exogenous story versions, we used different wording for the part of the story where the agent (X) is related to the object (Y). In the endogenous versions, we used formulations such as: “X is trying to find Y,” “X is trying to spot Y,” or “X is looking attentively for Y.” In the exogenous versions, we used formulations such as: “X’s eyes are drawn to Y,” “X’s gaze is captured by Y,” or “X’s attention is captured by Y.” We matched the average number of words across all four conditions (24 words). The number of stories that included the words “attention” or “attentively” was balanced between the endogenous and exogenous categories (43 stories in each). Among the 80 stories, for each subject, 20 were randomly selected to be used in the endogenous-self version; 20 in the endogenous-other version; 20 in the exogenous-self version; 20 in the exogenous-other version. Thus, for the example story shown in Figure 1B, each subject saw only one of the four versions. In this manner, each subject saw 80 social stories, 20 of each type, balanced for as many properties as possible other than the two factors that were manipulated.

Finally, we constructed 20 additional stories for the non-social control condition (Figure 1B). To keep the control stories as semantically similar as possible to the social stories, we based them on a subset of the 80 original stories. Crucially, the agent in the original story was substituted with a non-human object, such as a camera or a spotlight, that has a source and a target just as attention does. For instance, the original story, “You are in a bike shop, and numerous bikes hang on one of the walls. You are attentively looking for that red Italian sports bike,” was adapted to the non-social condition by substituting the agent with a spotlight: “In a bike shop, on one of the walls, hangs numerous bikes. A bright spotlight is shining on a red Italian sports bike.” The average number of words of the non-social stories (24 words, standard deviation = 3) was matched with the attention stories.

The purpose of the probe statement at the end of each trial was to ensure that subjects carefully read the stories. Each statement described one detail of the preceding story that could be either true or false. We restricted the probe statements to the spatial context of the story (place probe: e.g., “Emma is on a bus”) or the object being described (object probe: e.g., “The Van Gogh painting has sunflowers”) in order to avoid alerting subjects to the focus of the experiment on theory of mind and attention. Half of the probe statements were place probes and half object probes. Within both the place and the object probes, half were true and half were false. The probe was on screen for 4 s, during which subjects were required to indicate whether the statement was true or false by button press.

The experiment consisted of 10 runs of approximately 4 min each. In each run, the 5 conditions were repeated 2 times, yielding a total of 10 trials per run. The trial order was randomized, with the limitation that two consecutive trials could not belong to the same condition. Each run included 18 s of baseline before the onset of the first trial and 12 s of baseline after the offset of the last trial.

### Post-scan questionnaire

At the end of the scanning session, subjects were asked what they thought the 1 purpose of the experiment was and what they thought it was testing.

### fMRI data acquisition

Functional imaging data were collected using a Siemens Prisma 3T scanner equipped with a 64-channel head coil. Gradient-echo T2*-weighted echo-planar images (EPI) with blood-oxygen dependent (BOLD) contrast were used as an index of brain activity (Logothetis et al., 2001). Functional image volumes were composed of 54 near-axial slices with a thickness of 2.5 mm (with no interslice gap), which ensured that the entire brain excluding cerebellum was within the field-of-view in all subjects (54 x 78 matrix, 2.5 mm x 2.5 mm in-plane resolution, TE = 30 ms, flip angle = 80**°**). Simultaneous multi-slice (SMS) imaging was used (SMS factor = 2). One complete volume was collected every 2 s (TR = 2000 ms). A total of 1300 functional volumes were collected for each participant, divided into 10 runs (130 volumes per run). The first three volumes of each run were discarded to account for non-steady-state magnetization. A high-resolution structural image was acquired for each participant at the end of the experiment (3D MPRAGE sequence, voxel size = 1 mm isotropic, FOV = 256 mm, 176 slices, TR = 2300 ms, TE = 2.96 ms, TI = 1000 ms, flip angle = 9°, iPAT GRAPPA = 2). At the end of each scanning session, matching spin echo EPI pairs (anterior-to-posterior and posterior-to-anterior) were acquired for blip-up/blip-down field map correction.

### FMRI preprocessing

Results included in this manuscript come from preprocessing performed using FMRIPREP version 1.2.3 (Esteban et al., 2019) (RRID:SCR_016216), a Nipype (Gorgolewski et al., 2011) (RRID:SCR_002502) based tool. Each T1w (T1-weighted) volume was corrected for INU (intensity non-uniformity) using N4BiasFieldCorrection v2.1.0 (Tustison et al., 2010) and skull-stripped using antsBrainExtraction.sh v2.1.0 (using the OASIS template). Spatial normalization to the ICBM 152 Nonlinear Asymmetrical template version 2009c (Fonov et al., 2009) (RRID:SCR_008796) was performed through nonlinear registration with the antsRegistration tool of ANTs v2.1.0 (Avants et al., 2008) (RRID:SCR_004757), using brain-extracted versions of both T1w volume and template. Brain tissue segmentation of cerebrospinal fluid (CSF), white-matter (WM) and gray-matter (GM) was performed on the brain-extracted T1w using fast 10 (Zhang et al., 2001) (FSL v5.0.9, RRID:SCR_002823).

Functional data was slice time corrected using 3dTshift from AFNI v16.2.07 (Cox, 1996) (RRID:SCR_005927) and motion corrected using mcflirt (FSL v5.0.9) (Jenkinson et al., 2002). This was followed by co-registration to the corresponding T1w using boundary-based registration (Greve and Fischl, 2009) with six degrees of freedom, using flirt (FSL). Motion correcting transformations, BOLD-to-T1w transformation and T1w-to-template Montreal Neurological Institute (MNI) warp were concatenated and applied in a single step using antsApplyTransforms (ANTs v2.1.0) using Lanczos interpolation.

Many internal operations of FMRIPREP use Nilearn (Abraham et al., 2014) (RRID:SCR_001362]) principally within the BOLD-processing workflow. For more details of the pipeline see https://fmriprep.readthedocs.io/en/latest/workflows.html.

### Testing prediction 1

The purpose of the first analysis was to determine whether the brain encoded information concerning the type of attention (endogenous or exogenous) present in the stories. For this analysis, we used MVPA, which tests whether patterns of brain activity can be used to decode the distinction between two conditions. It is a more sensitive analysis than the more common, simple subtraction methods. The reason for using this sensitive measure is that the difference between exogenous and endogenous trial types was extremely subtle. Both trial types engaged social cognition, and therefore might cancel each other out in a simple subtraction. The stimuli were nearly identical, differing only in a few words that indicated the type of attention used by the agent in the story. In addition, the type of attention featured in the story was irrelevant to the task performed by the subject. To accommodate the subtlety of the distinction between conditions, we designed the study to use MVPA. We hypothesized that with MVPA, brain activity would carry information about the endogenous versus exogenous distinction; and that decoding would be evident in regions of interest (ROIs) within the network of areas typically found to be involved in social cognition, especially within the TPJ.

We defined our ROIs as spheres centered on the statistical peaks reported in an activation likelihood estimation (ALE) meta-analysis of 16 fMRI studies (including 291 subjects) involving theory-of-mind reasoning (van Veluw and Chance, 2014), in accordance with generally accepted guidelines in ROI analysis (Poldrack, 2007). The ROIs are shown in Figure 2. The peaks were located in six areas: the left TPJ (Montreal Neurological Institute [MNI]: -52, -56, 24), right TPJ (MNI: 55, -53, 24), left STS (MNI: 22 -59, -26, -9), right STS (MNI: 59, -18, -17), MPFC (MNI: 1, 58, 19), and the precuneus 23 (MNI: -3, -56, 37). The radius of the ROI spheres was 10 mm, corresponding to the approximate volume (4,000 mm^3^) of the largest clusters (TPJ and MPFC) reported in van Veluw and Chance (2014). The same sphere radius was used for all ROIs.

**Figure 2.**
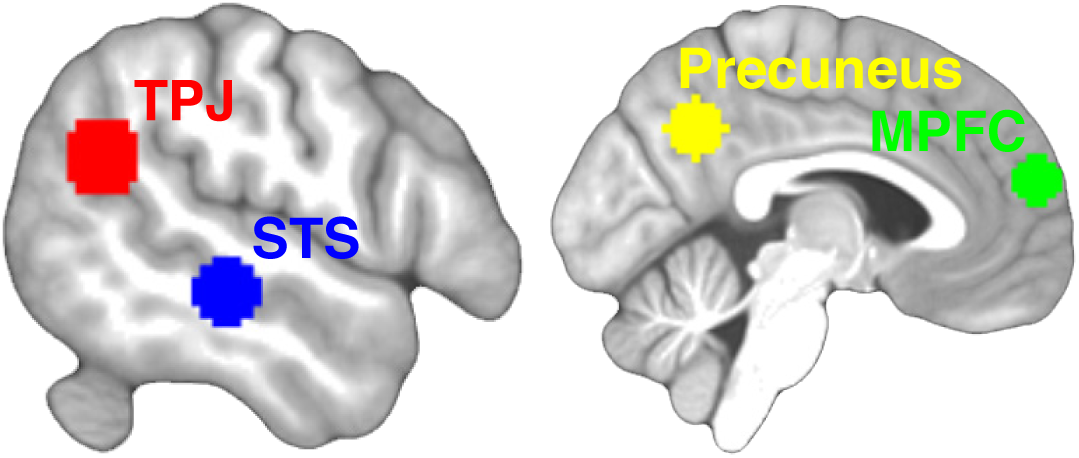
Regions of interest (ROIs). Six ROIs were defined based on peaks reported in an activation likelihood estimation meta-analysis of 16 fMRI studies involving theory-of-mind reasoning (van Veluw and Chance, 2014). The ROIs consisted of 10-mm-radius spheres centered on peaks in the bilateral temporoparietal junction (TPJ) and superior temporal sulcus (STS), and two midline structures: the precuneus and medial prefrontal cortex (MPFC). Here, the TPJ and STS ROIs on the left side are shown.

The fMRI data from all participants were analyzed with the Statistical Parametric Mapping software (SPM12) (Wellcome Department of Cognitive Neurology, London, UK) (Friston et al., 1994). We first used a conventional general linear model (GLM) to estimate regression (beta) coefficients for each individual trial (i.e., 100 regressors), focusing on the 10-s story presentation phase of each trial. One regressor of no interest modelled the 4-s probe statement phase across conditions. Each regressor was modeled with a boxcar function and convolved with the standard SPM12 hemodynamic response function. In addition, ten run-specific regressors controlling for baseline differences between runs, and six motion regressors, were included. The trialwise beta coefficients for the endogenous and exogenous conditions (i.e., 80 beta maps) were then submitted to subsequent multivariate analyses (Haxby et al., 2001).

The MVPA was carried out using The Decoding Toolbox (TDT) version 3.999 (Hebart et al., 2015) for SPM. For each subject and ROI, we used linear support vector machines (SVMs, with the fixed regularization parameter of C = 1) to compute decoding accuracies. To ensure independent training and testing data sets, we used leave-one-run-out cross-validation approach. For each fold, the betas across all training runs were normalized relative the mean and standard deviation, and the same Z-transformation was applied to the betas in the left-out test run (Misaki et al., 2010). An SVM was then trained to discriminate activity patterns belonging to the endogenous or exogenous trials in nine runs, and then tested on the left-out run, repeated for all runs, resulting in a run-average decoding accuracy for each ROI and subject.

For statistical inference, the true group mean decoding accuracy was compared to a null distribution of group mean accuracies obtained from permutation testing. The same MVPA was repeated within each subject and ROI using permuted condition labels (10,000 iterations). A p value was computed as (1+the number of permuted group accuracy values > true value)/(1+the total number of permutations). To control for multiple comparisons across the six ROIs, we used the false discovery rate (FDR) correction (Benjamini and Hochberg, 1995). In addition, we also computed a bootstrap distribution around the true group mean accuracy by resampling individual-subject mean accuracies with replacement (10,000 iterations), from which a 95% confidence interval (CI) was derived (Nakagawa and Cuthill, 2007). A corrected p value < 0.05 in combination with a 95% CI that does not cross chance level were interpreted as a significant decoding effect at the group level (Nakagawa and Cuthill, 2007).

In addition, as further exploratory statistics beyond the targeted hypotheses of this study, we used a whole-brain searchlight analysis (Kriegeskorte et al., 2006) to test for possible areas of decoding outside the ROIs. This searchlight analysis is described in the *Supplementary Information* (SI Text S1, Figures S1-S4, and Tables S1-S4).

### Testing prediction 2

The purpose of the second analysis was to determine whether the brain encoded information concerning the type of agent (self versus other) present in the stories. The analysis methods were the same as for testing hypothesis 1, except that for regressors of interest we used the self-related and other-related trials, collapsed across the type of attention (exogenous or endogenous). Just as for hypothesis 1, we tested the six defined ROIs within the theory-of-mind network.

### Testing prediction 3

The purpose of the third analysis was to test for an interaction between the two variables (endogenous versus exogenous, and self versus other). We used MVPA to test whether the decoding for the type of attention was significantly different between the self-related and the other-related stories. The analysis methods were similar to those used for testing hypothesis 1 and 2, except in the following ways. We computed two MVPA decoding results, the first for distinguishing endogenous-self from exogenous-self stories, the second for distinguishing endogenous-other from exogenous-other stories. We then computed the difference between the two decoding results ([endogenous-self versus exogenous-self] – [endogenous-other versus exogenous-other]) to create a decoding difference score. Just as for hypothesis 1 and 2, we tested the six defined ROIs within the theory-of-mind network(van Veluw and Chance, 2014).

### Testing prediction 4

The purpose of the fourth analysis was to confirm whether our story stimuli engaged social cognition and thereby recruited brain areas within the expected theory of mind network. The analysis was meant as an added control to check the validity of the paradigm. The analysis methods were similar to those used for testing hypothesis 1-3, except in the following ways. We computed four MVPA decoding results: endogenous-self versus nonsocial, endogenous-other versus nonsocial, exogenous-self versus nonsocial, and exogenous-other versus nonsocial. (Because using MVPA to compare two conditions requires equal numbers of trials in both conditions, it was not possible to use a single analysis to compare all 80 social trials to the 20 nonsocial trials.) Each analysis represents a separate, alternative way to assess the social-versus-nonsocial decoding. Just as for hypothesis 1-3, we tested the six defined ROIs within the theory-of-mind network (van Veluw and Chance, 2014).

### Eye tracking analysis

Eye movements were recorded via an MRI-compatible infrared eye tracker (SR Research EyeLink 1000 Plus), mounted just below the projector screen, sampling at 1000 Hz. Before each scanning session, a calibration routine on five screen locations was used and repeated until the maximum error for any point was less than 1°. The obtained eye position data was cleaned of artifacts related to blink events and smoothed using a 20-ms moving average. We then built an SVM decoding model analogue to the cross-validation approach used for the fMRI data, but here based purely on eye tracking data, to test whether eye movement dynamics alone were sufficient to decode the conditions of interest (endogenous versus exogenous, and self versus other). In keeping with a previous study (Schneider et al., 2013), we organized the data in the following way. The part of the display within which the stimuli appeared was divided into an 8 x 4 grid of 32 equally sized squares. The grid covered the screen area within which the stories were presented (see red outline in Figure S5), and approximately corresponded to the locations of individual words (four lines, with eight words per line). For each trial, the proportion of time that the subject fixated within each square (32 features) and the saccades between those regions (32 x 32 = 1024 features) was calculated. These 1056 features, representing information about both where people were looking as well as saccade dynamics, were then averaged across repetitions for each of the four main conditions within each of the 10 runs, yielding one eye movement feature vector per condition per run (per subject). The feature vectors were submitted to an SVM classifier (C = 1). Using a leave-one-run-out approach, the SVM model was trained on endogenous versus exogenous story types, and then tested in the left-out run. At the group level, the decoding accuracies were tested against chance level using t-tests. A similar analysis was then performed on the contrast between self-related stories versus other-related stories. The results showed that endogenous-versus-exogenous and self-versus-other story types could not be decoded significantly better than chance using the pattern of eye movement. See *Supplementary Information* (SI Text S2 and Figure S5) for the results of the eye-tracking analysis.

## Data availability

The data that support the findings of this study are available at https://figshare.com/s/c3463d15bc78106a1b5c.

## Results

### Prediction 1

We hypothesized that participants would encode the attentional state of the agents in the stories in enough detail to distinguish between endogenous and exogenous attention, even though the difference between the story types was extremely subtle – only a few words that very slightly altered the semantic meaning of the sentences. We made the strong prediction that decoding would be found within the set of brain areas typically included in the theory-of-mind cortical network. In particular, based on prior studies (Igelström et al., 2016; Kelly et al., 2014), we anticipated that the decoding would be most evident in the TPJ. Figure 2 shows six ROIs within the theory-of-mind network, based on a meta-analysis of previous theory-of-mind studies (van Veluw and Chance, 2014). Figure 3A shows the results (see Table 1 for numerical details). Decoding accuracy for endogenous versus exogenous stories was significantly above chance for the left TPJ, and the significance of the left TPJ decoding survived a multiple comparison correction for the six ROIs (mean decoding accuracy 52.9%, 95% CI 50.7 to 55.2, p_uncorrected_=0.0046, p_FDR-corrected_=0.0276). The results confirm our central prediction. The left TPJ showed significant decoding of the attentional state – exogenous versus endogenous – of agents in a story. (See *Supplementary Information* for the results of an exploratory, brain-wide, searchlight analysis.)

**Table 1.** Decoding attention type, agent, and the interaction between the two, within the six ROIs. For definition of ROIs, see Figure 2. Mean decoding accuracy (%), 95% confidence interval (based on bootstrap distribution), and p value (based on permutation testing) are shown for each of the six ROIs. Results shown for decoding endogenous (endo) versus exogenous (exo) attention type, self versus other agent type, and the interaction between the two variables. *significant p values that survived correction for multiple comparisons across all six ROIs (corrected p < 0.05).

|  |  | L TPJ | R TPJ | L STS | R STS | MPFC | Precuneus |
| --- | --- | --- | --- | --- | --- | --- | --- |
| **Endo vs. Exo** | Mean accuracy | 52.9% | 51.4% | 50.4% | 48.0% | 49.5% | 50.2% |
|  | 95% CI | 50.7 – 55.2 | 49.1 – 53.9 | 47.8 – 52.8 | 45.9 – 50.1 | 47.6 – 51.4 | 48.5 – 51.8 |
|  | P value | 0.0046* | 0.1148 | 0.3518 | 0.9547 | 0.6439 | 0.4428 |
| **Self vs. Other** | Mean accuracy | 53.0% | 51.0% | 52.3% | 51.3% | 52.6% | 52.7% |
|  | 95% CI | 50.1 – 55.6 | 48.5 – 53.4 | 50.6 – 54.1 | 48.9 – 54.1 | 50.5 – 55.0 | 50.4 – 55.0 |
|  | P value | 0.0053* | 0.1974 | 0.0204* | 0.1241 | 0.0105* | 0.0099* |
| **(Self vs. Other) × (Endo vs. Exo)** | Mean accuracy diff | 1.6% | 1.5% | 2.0% | -3.0% | 2.5% | 0.6% |
|  | 95% CI | -2.7 – 6.3 | -3.6 – 6.5 | -2.7 – 6.3 | -7.4 – 1.1 | -2.3 – 6.7 | -5.5 – 5.5 |
|  | P value | 0.2430 | 0.2639 | 0.1967 | 0.8944 | 0.1414 | 0.3900 |

**Figure 3.**
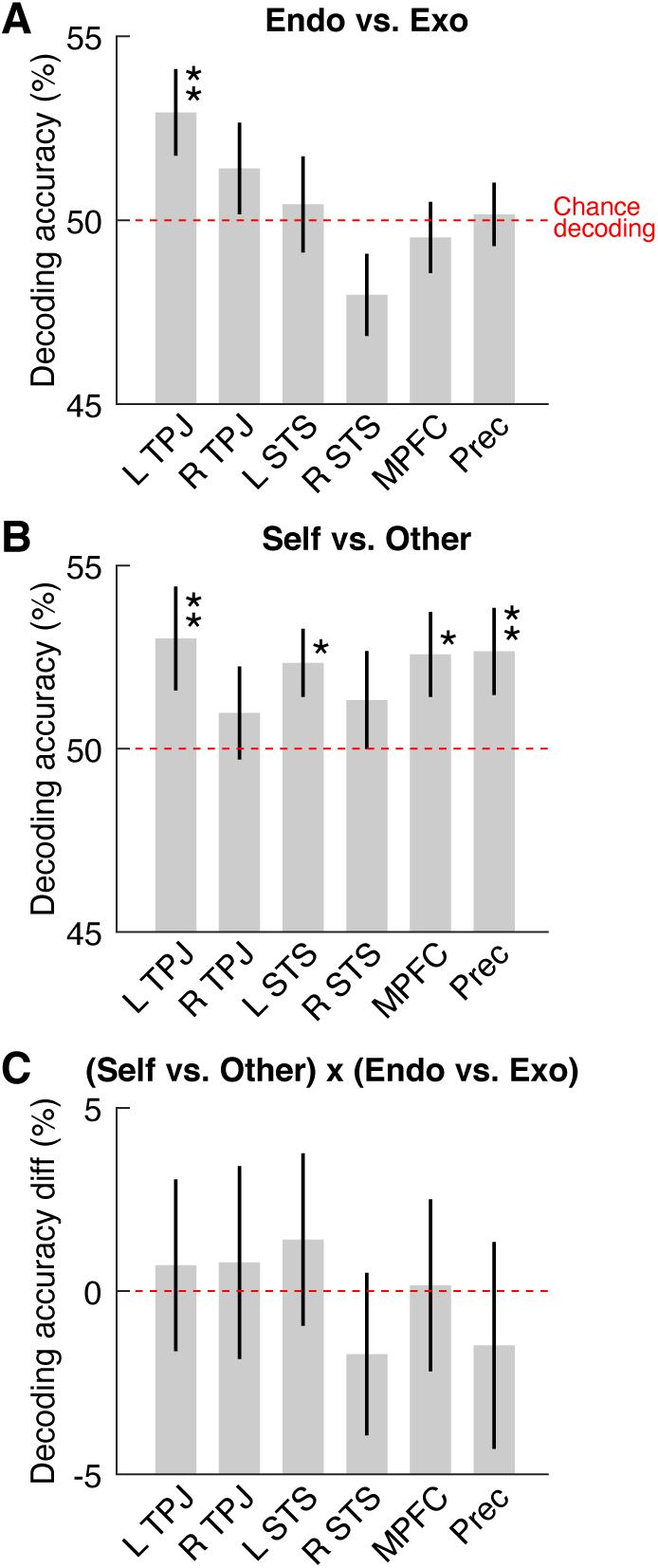
Decoding attention type, agent, and the interaction between them, in six brain areas. For definition of ROIs, see Figure 2. Each point shows mean decoding accuracy. Error bars show SEM. Red horizontal line indicates chance level decoding. Significance indicated by * (p<0.05) and ** (p<0.01), based on permutation testing (all significant p values also survived correction for multiple comparisons across all six ROIs [all corrected ps < 0.05]). **A**. The ability of a classifier, trained on BOLD activity patterns within each ROI, to decode endogenous (endo) versus exogenous (exo) attention. **B**. Decoding accuracy for agent (self versus other). **C**. Decoding accuracy for the interaction between type of attention and agent.

### Prediction 2

We hypothesized that participants would process the distinction between the two types of agent in the stories (self versus other). We made the strong prediction that decoding would be found within the same set of ROIs in the theory-of-mind cortical network. Figure 3B shows the results (see Table 1 for numerical details). Decoding accuracy for self versus other stories was significantly above chance, and survived a multiple comparisons correction, for the left TPJ (mean decoding accuracy 53.0%, 95% CI 50.1 to 55.6, p_uncorrected_=0.0053, p_FDR-corrected_=0.0210), left STS (mean decoding accuracy 52.3%, 95% CI 50.6 to 54.1, p_uncorrected_=0.0204, p_FDR-corrected_=0.0306), MPFC (mean decoding accuracy 52.6%, 95% CI 50.5 to 55.0, p_uncorrected_=0.0105, p_FDR-corrected_=0.0210), and precuneus (mean decoding accuracy 52.7%, 95% CI 50.4 to 55.0, p_uncorrected_=0.0099, p_FDR-corrected_=0.0210). These results confirm that the present paradigm, using stories that are subtly different from each other, can obtain social cognitive results that are consistent with previous findings.

### Prediction 3

We hypothesized that areas in the theory-of-mind network would not only encode the distinction between endogenous and exogenous attention, but do so to a significantly different extent in self-related stories than in other-related stories. However, the results showed no significant interaction in any of the ROIs (Figure 3C and Table 1). Thus, we found no support for prediction 3.

### Prediction 4

Finally, we asked whether the activity in the theory-of-mind network would distinguish between social stories and nonsocial stories. This final analysis served as a control to check the validity of the story stimuli and confirm that they engaged social cognition as expected. We expected a signal of much greater magnitude in this analysis than in the analyses described above. The reason is that, as noted above, the types of social stories differed from each other by only a few words, and were nearly identical in semantic content; thus any brain signal reflecting those differences is expected to be subtle. The distinction between social and nonsocial stories, however, was much greater semantically, and therefore the evidence of decoding in the brain is expected to be of greater magnitude. Figure 4 shows the results (see Table 2 for numerical details). The results are separated into six ROIs, and for each ROI, separated into four individual analyses, corresponding to each of the four main social conditions contrasted with the nonsocial control. Decoding accuracy was significantly greater than chance in almost all analyses across the six ROIs. The right STS showed the least consistent evidence of decoding. The TPJ bilaterally and the precuneus showed the most consistent evidence of decoding. These results show strong evidence of decoding of the social versus nonsocial stimuli in the known theory-of-mind, cortical network.

**Table 2.**
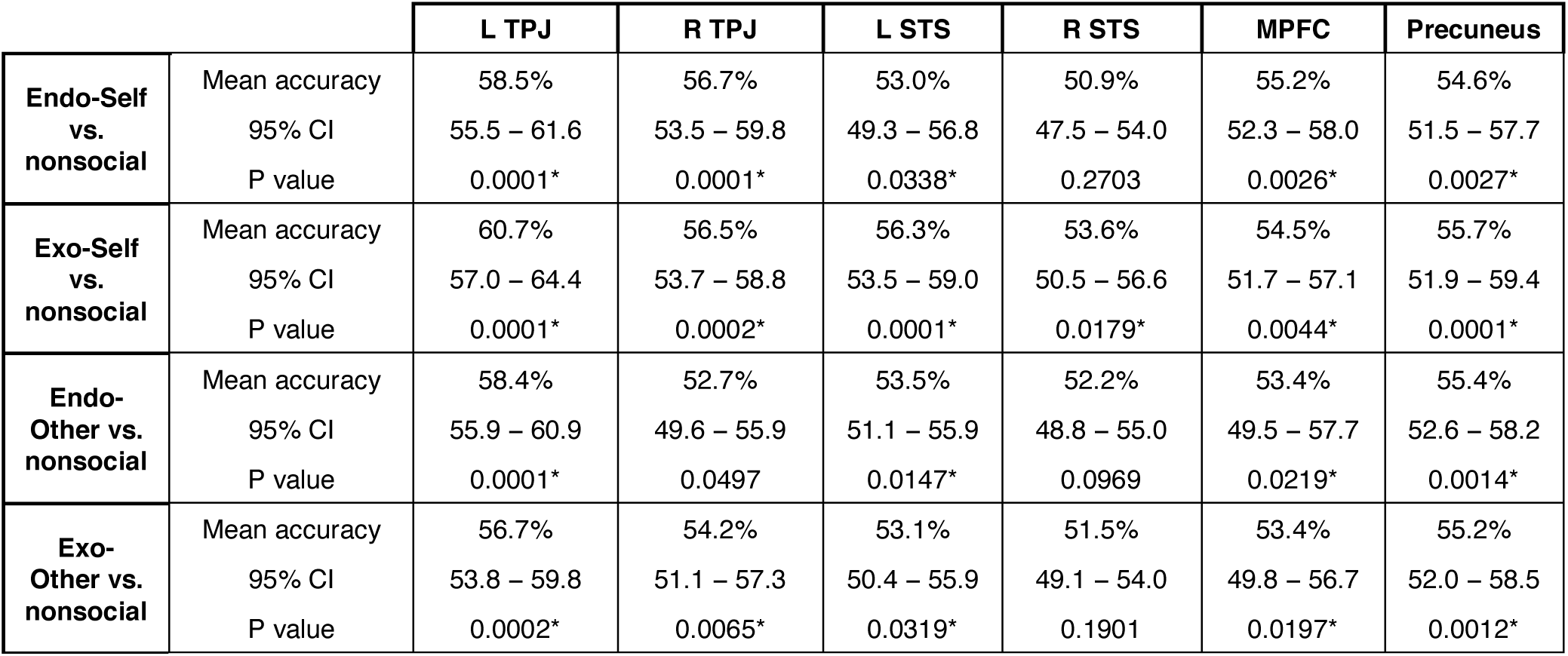
Decoding social versus nonsocial stories within the six ROIs. For definition of ROIs, see Figure 2. Mean decoding accuracy (%), 95% confidence interval (based on bootstrap distribution), and p value (based on permutation testing) are shown for each of the six ROIs. Results shown for each of four social story conditions (endogenous-self, exogenous-self, endogenous-other, and exogenous-other) versus the nonsocial baseline. *significant p values that survived correction for multiple comparisons across all six ROIs (corrected p < 0.05).

**Figure 4.**
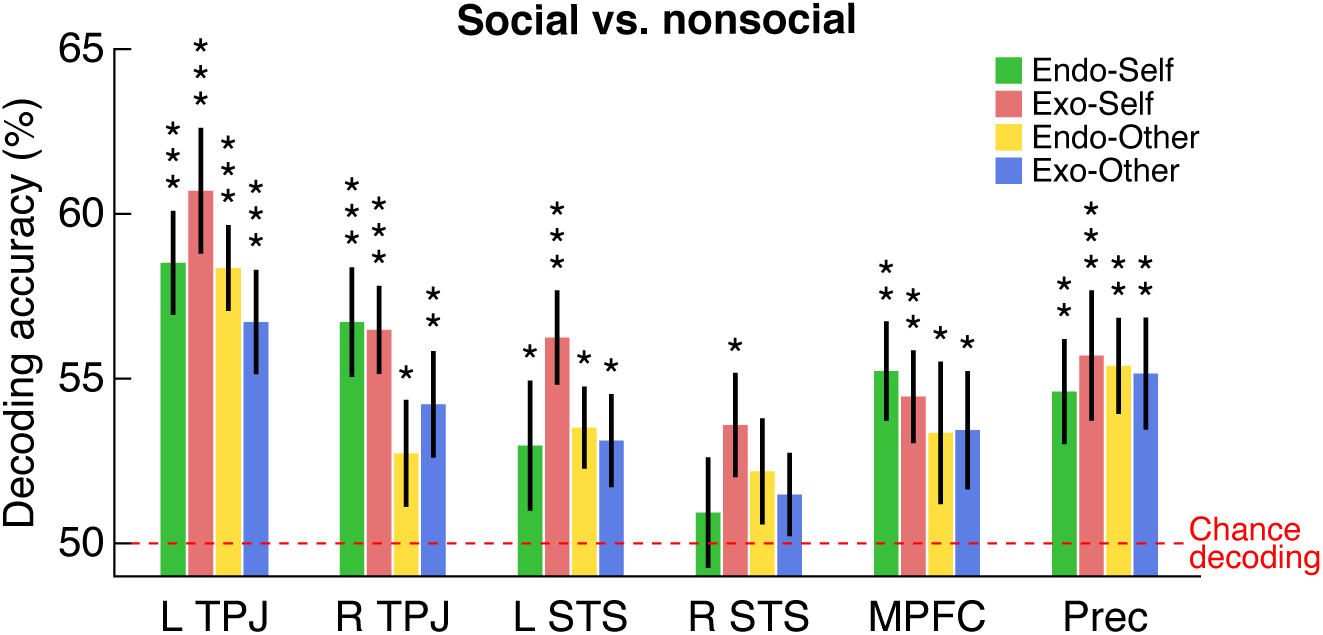
Decoding social versus nonsocial stories. The ability of a classifier, trained on BOLD activity patterns within each of the six ROIs, to decode each of the four social story conditions (endogenous-self, exogenous-self, endogenous-other, and exogenous-other) versus the nonsocial baseline. Each bar shows mean decoding accuracy, error bars show SEM, red horizontal line shows chance level decoding. Significance indicated by * (p<0.05), ** (p<0.01), and *** (p<0.001) based on permutation testing (all but one of the significant p values also survived correction for multiple comparisons across all six ROIs; see Table 2 for numerical details).

## Discussion

This study analyzed brain activity while people read stories about agents attending to objects in the environment. We examined whether specific brain areas could decode information about the type of attention referenced in the story (exogenous versus endogenous), and about the type of agent in the story (whether the agent was the subject reading the story or a different person). We hypothesized that if the brain constructs a model of attentional state that is used in social cognition, then areas of the brain known to be involved in social cognition should be able to distinguish between the two types of attention, exogenous and endogenous, represented in the stories. Our main analysis confirmed the hypothesis: the left TPJ showed significant decoding of information about endogenous versus exogenous attention. The finding is, arguably, remarkable, given that the semantic and wording difference between the two story types is extremely subtle.

These results support a new and growing body of evidence that the human brain constructs a model of attention to aid in theory of mind (Guterstam et al., 2019; Guterstam and Graziano, 2020; Kelly et al., 2014; Pesquita et al., 2016; Vernet et al., 2019). The model includes information about attention that is deeper and more complex than just gaze direction or an identification of the attended object. At least one aspect of attention incorporated into the model appears to be the manner in which attention moves to an object: endogenously (internally directed) or exogenously (externally induced). The processing of the model appears to engage the theory-of-mind cortical network. The left TPJ showed the strongest decoding result. It is not clear why the left hemisphere showed stronger activity than the right in the present task. Social cognition tasks often activate the TPJ bilaterally, but typically engage the right TPJ more (Saxe and Wexler, 2005). One speculation is that some aspect of the present task, perhaps explicitly instructing people that the task was a test of reading comprehension, caused an emphasis on linguistic processing, biasing the activity toward the left hemisphere. Other explanations for the left-hemisphere bias may also be possible.

We were also able to analyze brain areas involved in self-versus-other encoding. We found evidence of self-versus-other encoding in the left TPJ, left STS, MPFC and precuneus. The MPFC and precuneus have been previously implicated in self processing (Northoff et al., 2006; Ochsner et al., 2004; Passingham et al., 2010; Qin and Northoff, 2011; van Veluw and Chance, 2014), and the TPJ is consistently activated in fMRI studies involving self-recognition (van Veluw and Chance, 2014) and first-person perspective taking (Ionta et al., 2011). These results lend confidence to the present paradigm, showing that even the very subtle differences between our story stimuli were able to reveal cortical results consistent with previous studies.

Contrary to our prediction 3, we found no evidence for an interaction between attention type and agent type decoding in the theory-of-mind ROIs. (As noted in the *Supplementary Information*, during an exploratory searchlight analysis, we also found no evidence of an interaction effect in any other brain area.) Although it is possible that our paradigm was simply not sensitive enough to detect subtle interaction effects, these results suggest that the brain encodes information about attention type in a similar manner in the self and in others. Finally, significant decoding of the social-versus-nonsocial distinction was obtained across most of the theory-of-mind ROIs. This finding confirmed the validity of the paradigm, and was expected based on previous experiments of the theory-of-mind network (Gallagher et al., 2000; Saxe and Kanwisher, 2003; van Veluw and Chance, 13 2014; Vogeley et al., 2001).

The use of a story-reading paradigm allowed us to systematically manipulate the kind of attention represented in the stimulus while keeping other experimental factors close to identical. The endogenous and exogenous story versions differed only with respect to a few key words specifying the type of attention, while the rest of the stories were semantically the same. To avoid cognitive bias or expectation effects, the probe task performed by the subjects concerned details about the spatial context or the objects in the stories, effectively distracting subjects from the description of attention. A post-scan questionnaire confirmed that none of the subjects came close to figuring out the purpose of the experiment (which they had been told was a “Reading Comprehension Experiment”). The finding of brain areas that significantly decoded the type of attention, despite the distinction between endogenous and exogenous attention being subtle and task-irrelevant, indicates that the human brain automatically, and possibly also implicitly (Pesquita et al., 2016), constructs a model of an agents’ attention that specifies at least some dynamic aspects of how that attention is moving around the scene.

## Acknowledgments

This work was supported by the Princeton Neuroscience Institute Innovation Fund. Arvid Guterstam was supported by the Wenner-Gren Foundation, the Sweden-America Foundation, and the Promobilia Foundation. The authors would like to thank Sam Nastase for valuable input regarding the multivoxel pattern analysis.

## Competing interests

The authors declare no competing financial interests.

## Supplementary Information

### SI Text S1: Searchlight Analysis

Our primary analysis, described in the main text, was a targeted testing of strong hypotheses within a set of defined RIOs. Such targeted testing is preferred because it puts strong hypotheses to a direct test, and because it is more sensitive, avoiding the statistical problems of a brain-wide multiple comparison correction. Given the extremely subtle differences between the story types in the present experiment, such a targeted and sensitive analysis was preferred. However, in addition to the targeted ROI analyses, we also performed an exploratory analysis to ask whether any meaningful decoding activity might be identified outside of the ROIs. We used a searchlight analysis (Kriegeskorte et al., 2006). The searchlight analysis is fundamentally different from the ROI analysis. It is not targeted to specific brain areas on the basis of predictions. Instead, it is a whole-brain analysis that is much more statistically conservative because of the brain-wide multiple comparisons. In general, one would not expect the searchlight analysis to align perfectly with the ROI analysis. Activations revealed in the more sensitive ROI analysis might not appear in the searchlight analysis. Instead, it is exploratory in nature, and its usefulness is that it may reveal clusters of strong decoding in unanticipated areas outside the ROIs.

#### Endogenous-vs-exogenous searchlight analysis

First, the brain was partitioned into overlapping voxel clusters of spherical shape (10-mm radius). In each of these clusters, a decoding accuracy was computed using the same model input, SVM parameters, and procedures as described for the ROI analysis. This process resulted in an endogenous-versus-exogenous decoding accuracy map for each subject, in which the value of each voxel represents the average proportion of correctly classified trials relative to chance level (50%) based on the 10 mm sphere of tissue surrounding that voxel. The subject-wise decoding maps were then smoothed using a 3-mm full-width-half-maximum (FWHM) Gaussian kernel, and entered into a second-level analysis using SPM12. At the second-level, the whole-brain decoding maps were thresholded at p < 0.001 (uncorrected for multiple comparisons). For statistical inference, we employed a cluster-level, whole-brain approach to find clusters that passed the threshold of p < 0.05, corrected for brain-wide multiple comparisons using the familywise error rate correction as implemented by SPM12. In a purely descriptive manner, we also report strong decoding activity, defined as clusters ≥ 10 voxels using the cluster-forming threshold of p < 0.001 uncorrected, that did not survive correction at the whole-brain level (Table S1).

Clusters revealed in the searchlight analysis were projected onto orthogonal sections of the average structural scan generated from the 32 subjects for anatomical localization. The decoding clusters were also projected onto a 3D canonical brain surface using the software Surf Ice (University of South Carolina, McCausland Center for Brain Imaging). Figure S1 shows the brain-wide peak in decoding obtained with the searchlight method.

The searchlight analysis revealed no clusters that were brain-wide significant at the corrected p < 0.05 threshold. When examining the voxel-wise p < 0.001 threshold, four clusters (≥10 voxels) were observed (see Table S1 for details). The strongest peak decoding (t = 4.21) was located in the left posterior STS (within the TPJ; see Figure S1), thus consistent with the main, ROI analysis.

#### Self-versus-other searchlight analysis

The same method used for the exogenous-versus-endogenous searchlight analysis was used for the self-versus-other searchlight analysis. The analysis revealed four clusters that significantly decoded the self-versus-other distinction (p < 0.05) after correcting for multiple comparisons using the whole brain as search space. The global decoding peak was located on the left angular gyrus (part of TPJ; see Figure S2) (t = 5.92), which is compatible with our main ROI decoding results. All decoding clusters, including those passing the uncorrected threshold p < 0.001, are listed in Table S2.

#### Endogenous-versus-exogenous X self-versus-other searchlight analysis

The same method as in the previous sections was used for the interaction, or exogenous-versus-endogenous X self-versus-other, searchlight analysis. The analysis revealed no clusters that decoded the endogenous-versus-exogenous distinction significantly (p < 0.05, after correcting for multiple comparisons using the whole brain as search space) different in self-related compared to other-related stories. When examining the voxel-wise p < 0.001 threshold, two clusters were observed (see Table S3 for details). The strongest peak decoding (t = 4.20) was located in the left posterior STS (Figure S3). All decoding clusters, including those passing the uncorrected threshold p < 0.001, are listed in Table S3.

### Social-versus-nonsocial searchlight analysis

For this analysis, we performed four separate whole-brain searchlight analyses, testing for decoding that distinguished social from nonsocial stories. Each of the four analyses was restricted to comparing a specific type of social story to the nonsocial control: endogenous-self versus nonsocial, exogenous-self versus nonsocial, endogenous-other versus nonsocial, and exogenous-other versus nonsocial. We first identified clusters that passed a threshold of p < 0.05 corrected using the entire brain as search space. We then found brain areas of overlap, that were ≥10 voxels in size, between the decoding clusters obtained in the four different analyses. These areas of overlap represent brain regions that showed significant decoding for the social-versus-nonsocial comparison, in a consistent manner, across all types of social stimuli. Using this method, four brain areas were obtained (see Table S4 and Figure S4 for details).

### SI Text S2: Eye-Tracking Decoding

To examine whether eye movement dynamics could explain our fMRI decoding results, we systematically organized eye position and saccade data into gaze pattern vectors, and submitted it to a decoding model analogue to the one used for the fMRI data (see Methods for details). Out of 32 subjects, 27 had usable eye tracking data available (five subjects had data with unacceptable levels of noise due to either the presence of glasses or a partially occluded pupil). The results showed that this classifier, based solely on eye tracking data, could not decode attention type (endogenous-versus-exogenous decoding accuracy 55.2%, p=0.102) or agent (self-versus-other decoding accuracy 52.4%, p=0.376) significantly better than chance level. See Figure S5.

**Figure S1.**
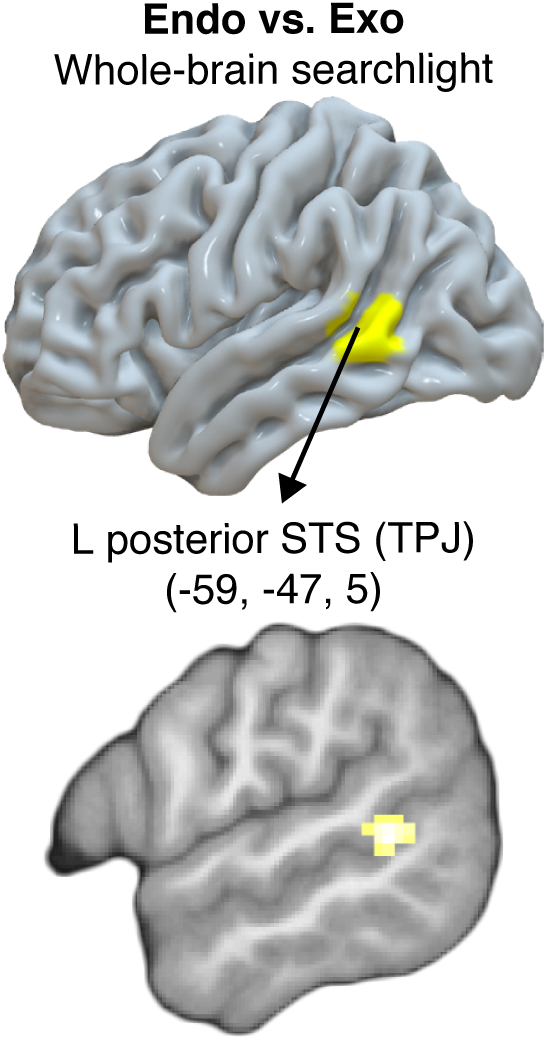
Decoding attention type at the whole-brain level. The cluster shown had the highest decoding accuracy in the whole-brain, searchlight analysis, for the endogenous-versus-exogenous comparison. See Table S1 for numerical details. Top: projected onto a 3D canonical brain surface. Bottom: projected onto a parasagittal section of the average structural scan generated from the 32 subjects for anatomical localization. For display purposes, the statistical threshold for the activation maps was set to p < 0.001, uncorrected.

**Figure S2.**
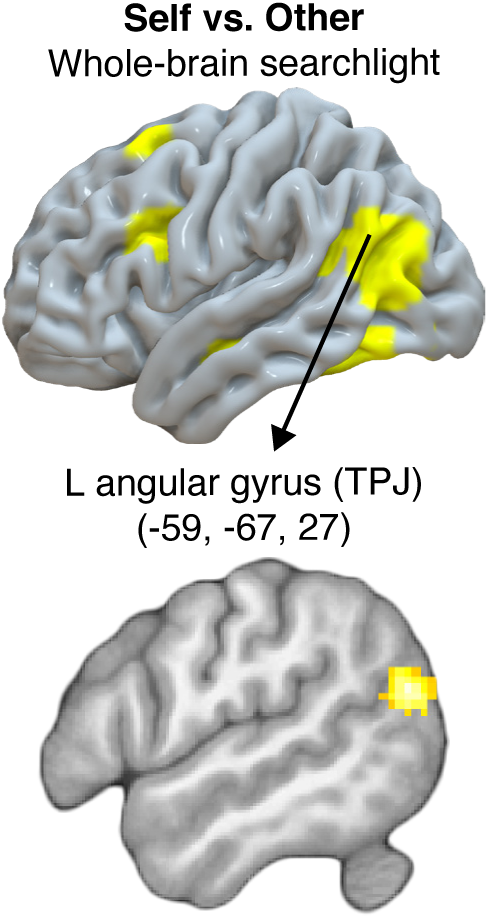
Decoding agent type at the whole-brain level. The cluster shown had the highest decoding accuracy in the whole-brain, searchlight analysis, for the self-versus-other comparison. See Table S2 for numerical details. Top: projected onto a 3D canonical brain surface. Bottom: projected onto a parasagittal section of the average structural scan generated from the 32 subjects for anatomical localization. For display purposes, the statistical threshold for the activation maps was set to p < 0.001, uncorrected.

**Figure S3.**
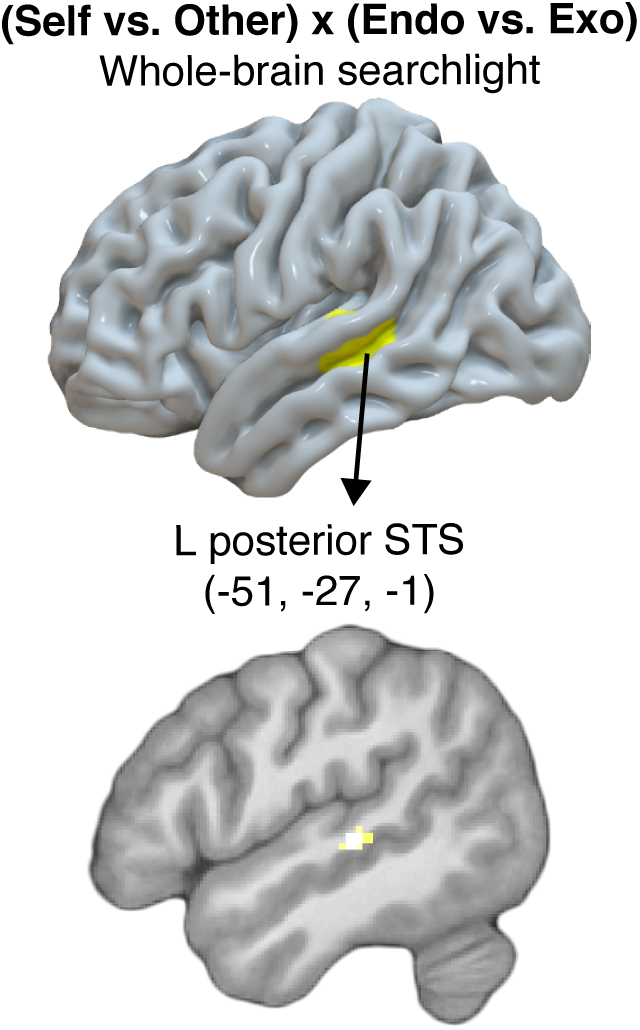
Decoding attention-by-agent interaction at the whole-brain level. The cluster shown had the highest decoding accuracy difference in the whole-brain, searchlight analysis, for the (endogenous-versus-exogenous)_SELF_ versus (endogenous-versus-exogenous)_OTHER_ comparison. See Table S3 for numerical details. Top: projected onto a 3D canonical brain surface. Bottom: projected onto a parasagittal section of the average structural scan generated from the 32 subjects for anatomical localization. For display purposes, the statistical threshold for the activation maps was set to p < 0.001, uncorrected.

**Figure S4.**
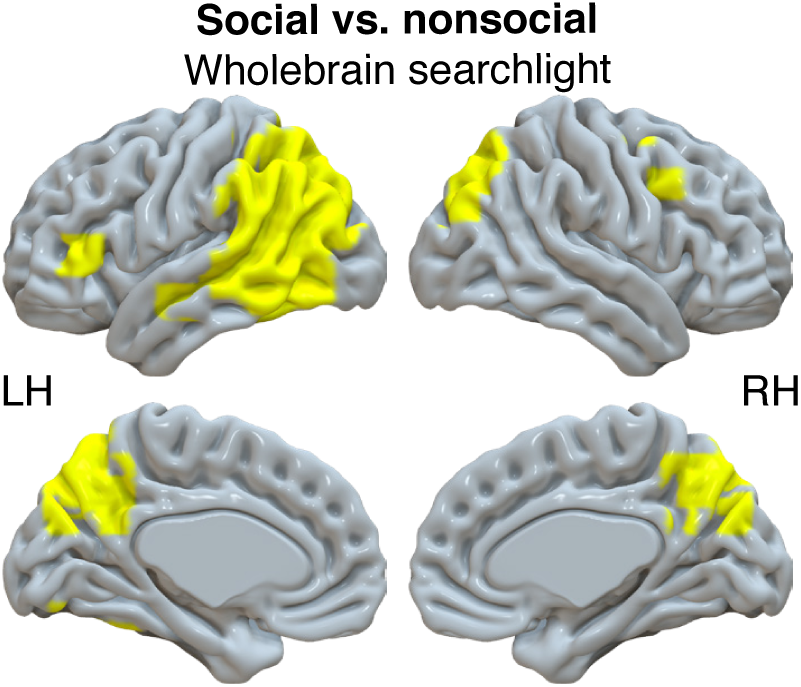
Decoding social versus nonsocial stories at the whole-brain level. The clusters shown are areas of overlap between four whole-brain, searchlight analyses, for the social-versus-nonsocial comparison (endogenous-self versus nonsocial, exogenous-self versus nonsocial, endogenous-other versus nonsocial, exogenous-other versus nonsocial). See Table S4 for numerical details. Projected onto a 3D canonical brain surface.

**Figure S5.**
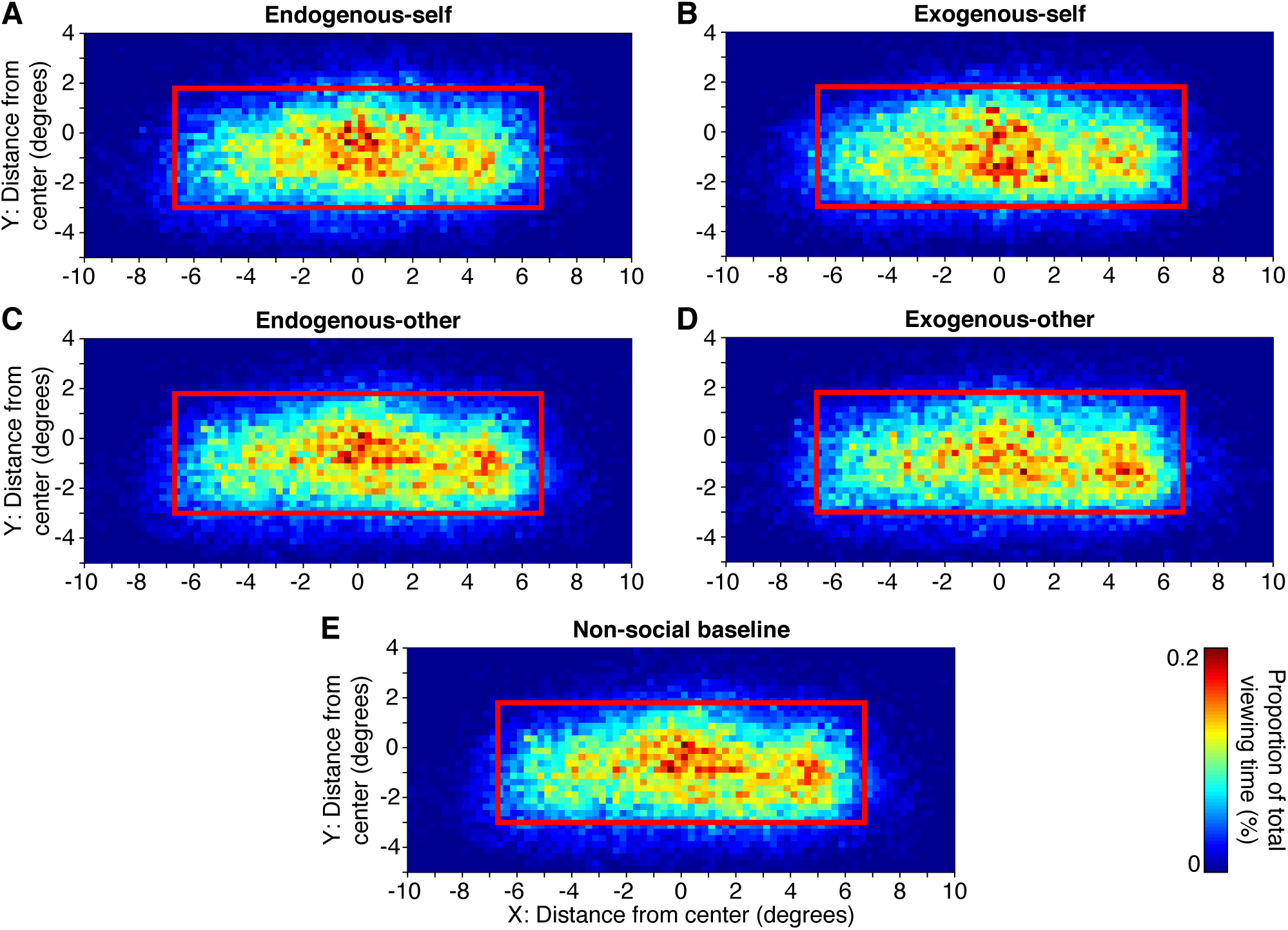
Eye tracking results. Subjects tended to fixate in a similar spatial pattern across the screen, and engage in similar saccade dynamics, regardless of the type of story presented. No significant ability to decode the story type based on the pattern of eye movement was obtained. The red rectangle in the heat maps in panels A-E indicates the area of the screen within which the story text appeared.

**Table S1.**
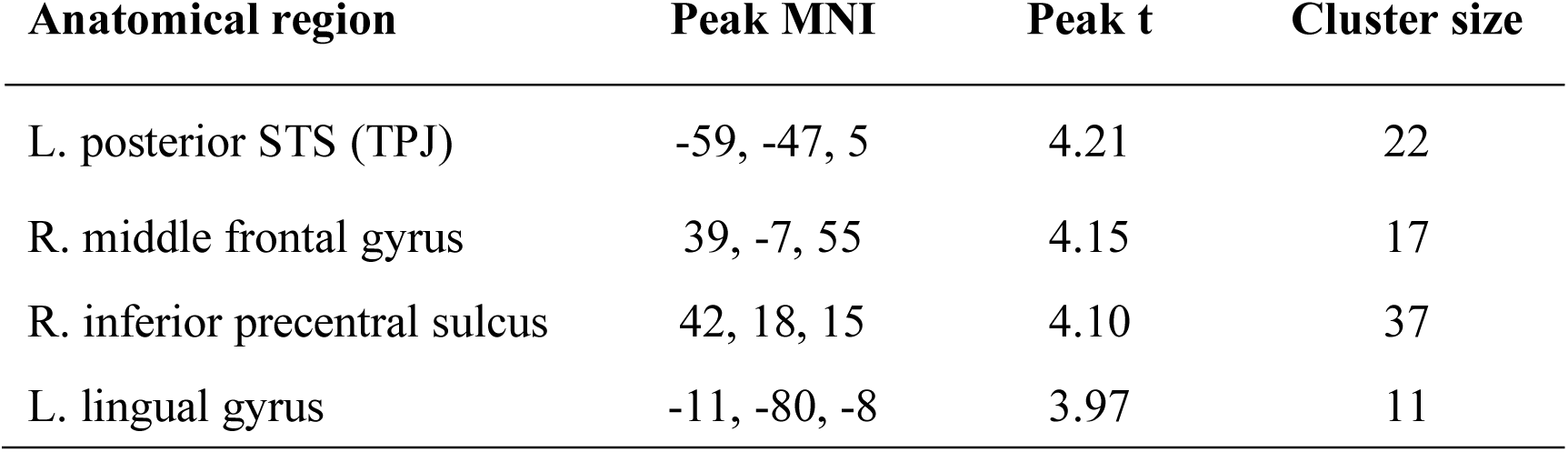
Decoding endogenous versus exogenous at the whole-brain level. All clusters (≥10 voxels) of decoding activity passing the voxelwise threshold of p < 0.001 (none of the clusters survived p < 0.05 correction for multiple comparisons using the whole brain as search space).

| Anatomical region | Peak MNI | Peak t | Cluster size |
| --- | --- | --- | --- |
| L. posterior STS (TPJ) | -59, -47, 5 | 4.21 | 22 |
| R. middle frontal gyrus | 39, -7, 55 | 4.15 | 17 |
| R. inferior precentral sulcus | 42, 18, 15 | 4.10 | 37 |
| L. lingual gyrus | -11, -80, -8 | 3.97 | 11 |

**Table S2.** Decoding agent type at the whole-brain level. All clusters (≥10 voxels) decoding agent type (self versus other) activity at the threshold of p < 0.001 (uncorrected). Corrected p values represent cluster-level correction using the whole brain as search space.

| Anatomical region | Peak MNI | Peak t | Cluster size | Corrected p value |
| --- | --- | --- | --- | --- |
| L. angular gyrus (TPJ) | -59, -67, 27 | 5.92 | 211 | <0.001 |
| R. precuneus | 4, -55, 30 | 5.91 | 57 | - |
| L. inferior temporal gyrus | -41, -15, -18 | 5.49 | 131 | 0.004 |
| L. inferior occipital gyrus | -31, -77, -8 | 5.26 | 54 | - |
| R. middle cingulate cortex | 2, -15, 40 | 4.24 | 47 | - |
| R. lingual gyrus | 24, -82, -8 | 4.82 | 121 | 0.006 |
| L. fusiform gyrus | -39, -57, -23 | 4.81 | 114 | 0.008 |
| R. calcarine sulcus | 7, -85, 5 | 4.71 | 53 | - |
| R. fusiform gyrus | 27, -37, -21 | 4.32 | 34 | - |
| R. inferior occipital gyrus | 47, -80, 3 | 4.05 | 27 | - |
| L. inferior frontal sulcus | -41, 13, 27 | 4.02 | 13 | - |
| L. superior frontal gyrus | -29, 16, 65 | 3.98 | 12 | - |

**Table S3.** Decoding attention-by-agent interaction at the whole-brain level. All clusters (≥10 voxels) in which endogenous-versus-exogenous decoding was better in self-related compared to other-related stories at p < 0.001 (uncorrected). (none of the clusters survived correction for multiple comparisons using the whole brain as search space).

| Anatomical region | Peak MNI | Peak t | Cluster size |
| --- | --- | --- | --- |
| L. posterior STS | -51, -27, -1 | 4.20 | 13 |
| R. inferior temporal gyrus | 47, -22, -28 | 3.96 | 13 |

**Table S4.** Decoding social versus nonsocial stories at the whole-brain level. Clusters (≥10 voxels) decoding social versus nonsocial stories significantly better than chance. The listed clusters represent the overlap of significant clusters (p < 0.05, corrected using a cluster-defining uncorrected threshold of p < 0.001 and the entire brain as search space) across four separate whole-brain searchlight analyses: endogenous-self versus nonsocial, exogenous-self versus nonsocial, endogenous-other versus nonsocial, and exogenous-other versus nonsocial.

| Anatomical region | Cluster size |
| --- | --- |
| L. TPJ | 1722 |
| L. and R. precuneus | 80 |
| R. intraparietal sulcus | 137 |
| L. inferior frontal sulcus | 22 |

#### SI Data S1. Story stimuli

List of all of the stories and story versions (endogenous-self, exogenous-self, endogenous-other, and exogenous-other) presented to the participants during the experiment.

